# Identification of RNA-based cell-type markers for the stem cell manufacture system with a statistical scoring function

**DOI:** 10.1101/2022.10.21.513209

**Authors:** Yu Shi, Weilong Yang, Haishuang Lin, Li Han, Alyssa J. Cai, Ravi Saraf, Yuguo Lei, Chi Zhang

**Author notes:** To whom correspondence should be addressed. (C.Z.); (C.Z.), (Y.L.). Department of Chemical Biology, School of Pharmaceutical Sciences, Peking University, Beijing 100191, China (Haishuang Lin).

## Abstract

Cell-type biomarkers are useful in stem cell manufacture to monitor cell purification, cell quantity, and quality. However, the study on cell-type markers, specifically for stem cell manufacture, is limited. The emerging questions are which RNA transcripts can serve as biomarkers during stem cell culture, and what method can efficiently and accurately discover these biomarkers. We developed a scoring function system to identify RNA biomarkers with RNA-seq data. We applied the method to two data sets, one for extracellular RNAs (ex-RNAs) and the other for intracellular microRNAs (miRNAs). The data set have RNA-seq data of ex-RNAs from cell culture media for six different types of cells, including human embryonic stem cells. To get the RNA-seq data from intracellular miRNAs, we cultured three types of cells: human embryonic stem cells (H9), neural stem cells (NSC), hESC-derived endothelial cells (EC) and conducted small RNA-seq to their intracellular miRNAs. Using these data, we identified a set of ex-RNAs/smRNAs as candidates of biomarkers for different types of cells for cell manufacture. We also used deep-learning based prediction methods and simulated data to validate these discovered biomarkers.

## Introduction

Within multicellular organisms, multiple cell types work together to carry out and characterize the normal functioning of different organs and tissues. Recognition of specific cell types is not only important for understanding the normal functions of cells and tissues, but also for studies on diseases, the development of therapeutic methods, and stem cell manufacture. Induced pluripotent stem cells (iPSCs) generated by reprogramming tissue cells can model diseases in vitro and treat diseases in vivo without significant immune rejection ^1,2^. Cell-type biomarkers are useful in stem cell manufacture to monitor cell purification, cell quantity, and quality. The purity is important because, in iPSC-derived regenerative medicine products, residual undifferentiated cells have a chance to form tumors after transplantation ^3,4^. Using an integrated in-line sensor, automated real-time monitoring of cell differentiation for cell manufacture also requires accurate cell-type biomarkers.

Most of the existing quality control tests are using biopsy, which involves crushing the cells and causes the cells that will be transplanted to be unable to be tested directly. Some biomarkers were used for detecting residual undifferentiated stem cells, such as LIN28A^4^ or miRNA-302 family ^5^. On the other hand, it has the potential to use extracellular RNAs (ex-RNAs) as biomarkers, a less invasive method than biopsy, to monitor cell purification, cell quantity, and quality. Cells secrete RNAs such as microRNAs (miRNAs), piwi-interacting RNAs (piRNAs), small nucleolar RNAs (snoRNAs), small nuclear RNAs (snRNAs), mRNA fragments, transfer RNA halves (tRHs), transfer RNA fragments (tRFs), and ribosomal RNA fragments (rRFs) as messengers to communicate and modulate other cells^6-8^. Ex-RNAs have been identified in biofluids (e.g., plasma) for various diseases ^9-12^ and are candidates for developing “Liquid Biopsy” technologies ^13^. The secreted RNAs (ex-RNAs) in cell culture medium have been reported ^9,14-19^. Studies show that smRNAs (∼15 to 200 nucleotides) in the culture media (ex-RNAs) can accurately characterize the heterogeneous cell population^14,15,17-19^. It will be challenging to find ex-RNAs that are highly expressed only in the target cell type with low/no expression in other 200 plus cell types. Fortunately, a cell biomanufacturing process only contains a limited number of cell types (e.g., 4 types). Literature shows when only having a few cell types, 20 plus cell-specific ex-RNA markers can be robustly identified ^9,17^. The emerging questions are what method can efficiently and accurately discover cell-type biomarkers from intracellular miRNAs or ex-RNAs for a cellular culture and cell differentiation system.

In this work, we developed a method to identify cell-type biomarkers from RNA-seq data for intracellular small RNAs or extracellular RNAs for a system with a limited number of cell types. Because our goal is to identify cell-type biomarkers for the automated real-time monitoring of cell differentiation for cell manufacture, we need to find candidate marker genes that have high expression in one cell type but low/no expression in the other few cell types. For a stem cell manufacturing system, the total number of cell types is small, and it could be two, three, or four different types of cells. The biomarkers that the method discovered are system-specific, and not necessary to be universal. For intracellular smRNAs, we cultured three types of cells: Human embryonic stem cell (H9), iPSCs-derived Neural stem cells (NSC), and iPSCs-derived Vascular endothelial cells (EC) and conducted smRNA-seq to their intracellular small RNAs. For extracellular RNAs in cell culture medium, we collected RNA-seq data for six different types of cells: Human embryonic stem cells (hESC) ^9^, human extrahepatic bile duct carcinoma cell line (KMBC) ^9^, peripheral blood B lymphocytes (B-cell) ^20^, Epidermal primary keratinocytes (PKCs) ^21^, lung cancer cell lines (A549)^22^, and colorectal cancer DKO cell line (DKO) ^23^. Using these datasets, we designed a scoring function to identify a set of smRNAs or ex-RNAs as candidates for biomarkers for cell-type prediction for cell manufacture. We synthesized gene expression data for different cell types and validated these discovered biomarkers with a machine learning model.

## Results

### Some smRNAs have high expression levels uniquely in the target cell type

As a cell-type-specific biomarker in a bioprocess, the expression level of the smRNA should have a stable and high expression level in the target cell type and close to zero expression level in other cell types. To develop computational methods to identify cell-type-specific markers accurately and efficiently, we collected data from two systems. First, we prepared H9 cells, a kind of human pluripotent stem cells, iPSCs-derived neural stem cells (NSCs), and iPSCs-derived endothelial cells (ECs) following our published protocols with three replicates from different batches ^24-27^. We conducted RNA-seq to smRNAs from H9, NSC, and EC. For each type of cell, three replicates from different batches were collected for sequencing. On average, each sample has 4 million cleaned short reads, and 400 miRNAs have read counts ≥ 5. After trimming, mapping, and counting, all reads from small RNA-seq for nine samples were normalized to reads per million total reads (RPM). Second, we collected transcriptomic data for extracellular RNAs (ex-RNAs) in cell culture media of six different types of cells: hESC, KMBC, B-cell, PKC, A549, and DKO^9,20-23^.

To test if the potential biomarker exists, we increased the cutoff of ex-RNA/smRNA abundance levels in the target cell type and decreased the ceiling of smRNA abundance levels in other cell types to see the number of remaining smRNAs. Figure 1A shows the result for ex-RNA dataset, when the target cell type was hESC. The bottom threshold for ex-RNA express level in hESC cells increased from 30 to 150 RPM. The total number of ex-RNAs decreased when the cutoff of ex-RNA express level in hESC was increased. For the other five types of cells, i.e. non-target cell type, an ex-RNA was kept if its expression level was smaller than the cutoff (ceiling cutoff), which was reduced as steps 20, 10, 5, and 2. If requiring ex-RNA abundance level in the target cell type, i.e. hESC, > 150 and that in other non-target types of cells < 2, 10 ex-RNAs were left. These ex-RNAs are candidates for cell-type markers for hESC cells because they have high expression levels in hESC (>>150 RPM), and low expression levels in any other five types of cells (<2 RPM). We got similar results if KMBC (Figure 1B), B-cell (Figure 1C), PKC (Figure 1D), A549 (Figure 1E), or DKO (Figure 1F) is considered as target cell types, respectively. There are 6, 8, 6, 3, and 3 candidate markers for cell-types KMBC, B-cell, PKC, A549, and DKO, respectively. For cell types hESC, KMBC, B-cell, PKC, and A549, the ceiling cutoffs for non-target cell types are stringent – the lowest cutoff is two or three. However, for DKO, the ceiling cutoffs for non-target cell types is 30. The total number of reads for DKO samples was much smaller than that for other samples. This leads to a difficulty to find a high-quality cell-type marker for DKO. Figure 1G-I show the result for the intracellular smRNA dataset. There are 2, 2, and 15 candidate markers for cell-type H9, NSC, and EC. For NSC cells, the bottom cutoffs and ceiling cutoffs are the loosest. This indicated that it is more difficult to find a good cell-type marker for NSC than for H9 or EC. These results indicate it is possible to find cell-type markers from smRNAs for a cell-manufacturing system, which has only several different types of cells, but a scoring function to identify the optimal markers in all candidates is required. From the distributions of ex-RNAs/smRNAs at different cutoffs, one can find that a cell-type-specific marker exists in a system of several different types of cells, but the number of this type markers is limited. On the contrary, the number of cell-type-nonspecific ex-RNAs or smRNAs is large. For example, ex-RNAs from *RNY1, RNAy4P7*, and *RNU5E*, exist in cell culture media for all six cell types with high abundances (>1000 RPM). For intracellular smRNAs, mir-302A, mir-302B, and mir-148A etc. have high abundances in all three different cell types.

**Figure 1.**
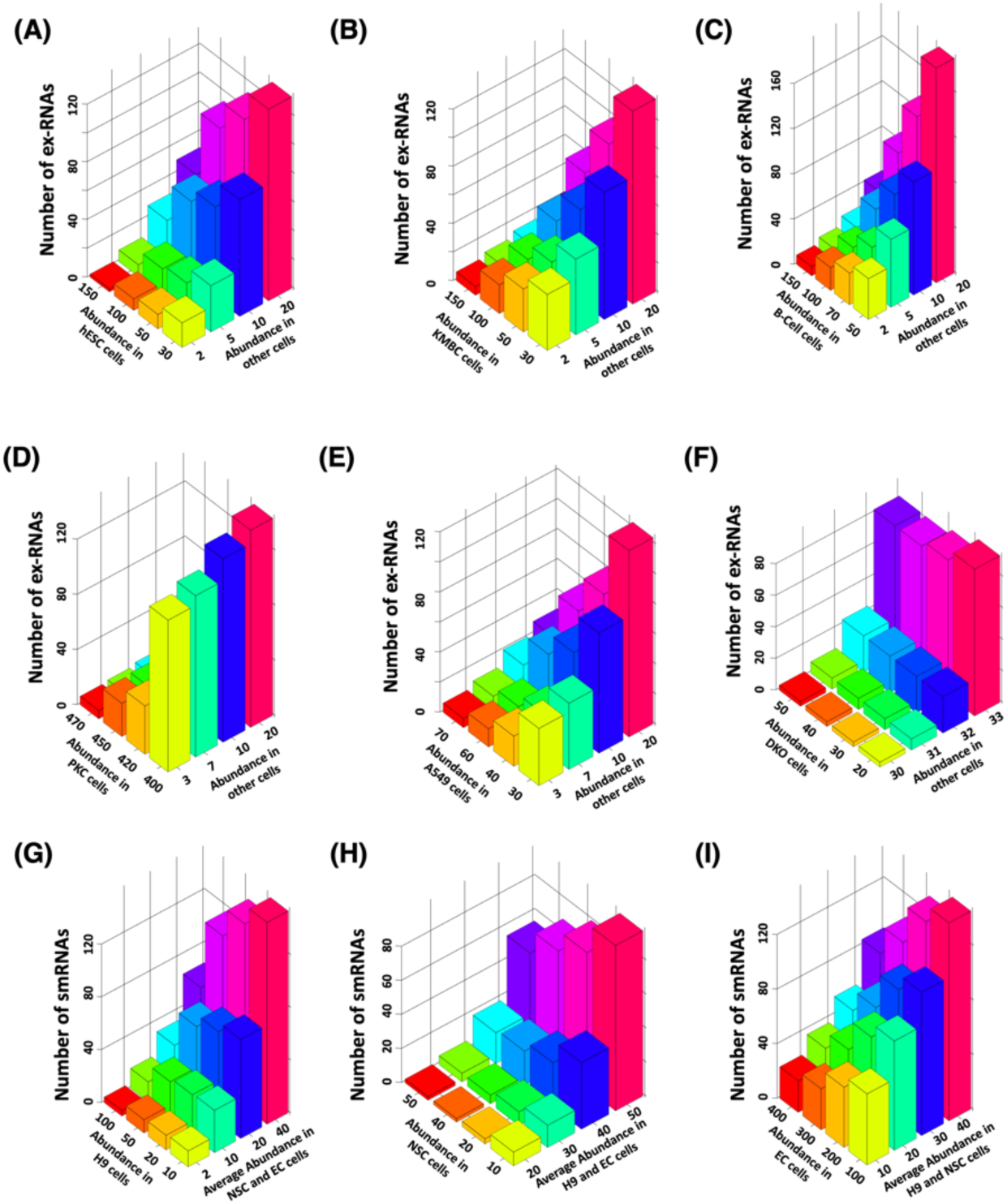
the numbers of smRNAs that are left for meeting abundance cutoffs. (A) the target cell type is hESC. (B) the target cell type is KMBC. (C) the target cell type is B-cell. (D) the target cell type is PKC. (E) the target cell type is A549. (F) the target cell type is DKO. (G) the target cell type is H9. (H) the target cell type is NSC. (I) the target cell type is EC. The X-axis is the bottom cutoff of ex-RNA/smRNA abundance in the target type of cells and the Y-axis is the ceiling cutoff of average ex-RNA/smRNA abundance in the any other non-target cell types.

### Identification of the cell type markers with the scoring function

An RNA-based cell-type marker shall have a high and stable expression level in the target cell but have a close-to-zero expression level in other types of cells. Therefore, we designed a statistical scoring function to rank RNAs transcribed from all genes. For one type of RNA, a score, S, was designed to describe its quality to serve as a marker for cell type. The scoring function is shown in Equation (1)

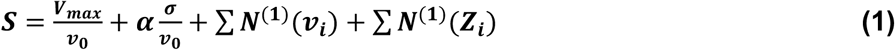

where *v*_*0*_ and σ are the mean expression level and standard deviation of the given RNA in all replicates for the target cell type, alpha is the imbalance score for replicates, *V*_*max*_ is the maximum of all RNA expression levels in the target cell type, *v*_*i*_ and *Z*_*i*_ are the mean expression level and its maximal Z-score in all replicates of the given RNA in the *i*-th of other cell types. *N*^(1)^ is the function for normalization. If *v*_*i*_ or *Z*_*i*_ is close to zero, *N*^(1)^ is recommended to be an exponential function. However, if *v*_*i*_ or *Z*_*i*_ is larger than zero, it is better to be divided by their maximal values in all RNAs.

From this scoring function, one can find that, for the target cells, the higher the ex-RNA/smRNA abundance levels the better, and the standard deviation of all replicates need to be small. Contrarily, for other non-target types of cells, the ex-RNA/smRNA abundance levels, and the maximal z-score for all three replicates are included to evaluate their low expression level and low fluctuations. The score of an ex-RNA/smRNA is the sum of these parts; It should be a positive number. For a good candidate for the cell-type marker, its score is the smaller the better.

We applied this scoring function to the dataset for ex-RNAs in the cell culture media of six cell types, hESC, KMBC, B-cell, PKC, A549, and DKO. The top 15 scores for these six cell types as the target cell are shown in Figure 2(A). For all six cell types, the first three scores are obviously smaller than other scores, indicating the top-ranked three ex-RNAs are good candidates of cell-type markers. These top-ranked cell-specific ex-RNA markers were identified, and their expression levels are shown in the heatmap Figure 2(B). For each cell type, the top-three ex-RNAs were selected as the candidates for cell-type markers because top-3 genes have relatively small scores in all six types of cells. For example, RNAs from *MTRNR2L2, TMEM215, MT-TN:Mt_tRNA* was ranked as the top ones for hESC cells. For the intracellular smRNAs, Figure 2(C) shows the top 9 scores for H9, NSC, and EC as the target cells, respectively. For each cell type, top-2 smRNAs were selected as the candidates for cell-type markers because top-2 genes have relatively small scores in all three types of cells. For H9 cells, miR-518b and miR-518c were selected as the candidate cell-type markers. MiR-3663 and miR-548ah were selected for NSC cells while VTRNA1-2 and THBS1 were for EC cells. The expression levels of these intracellular smRNAs are shown in Figure 2(D).

**Figure 2.**
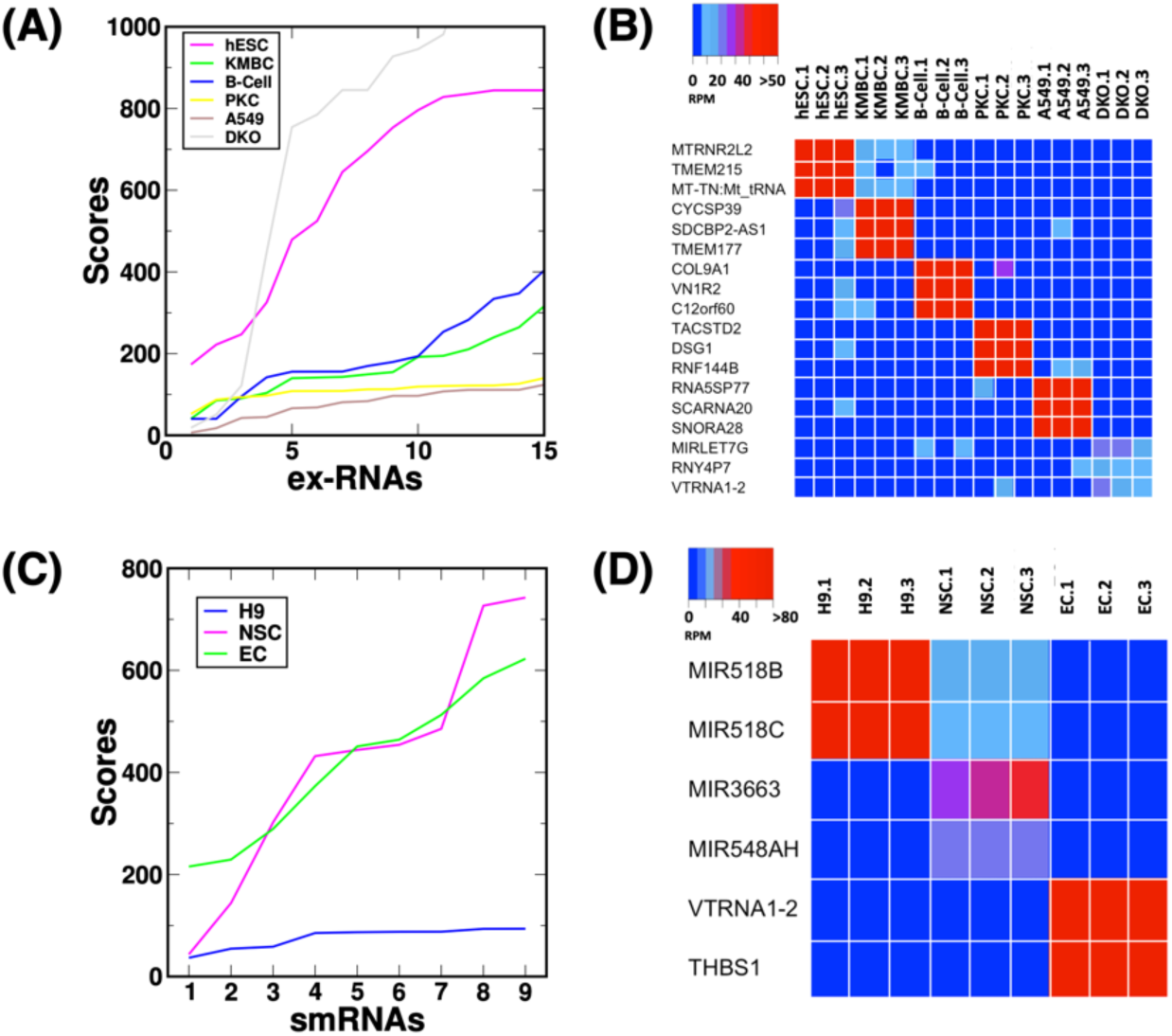
Result of the application of scoring function to real datasets. (A) the scores of the top 20 ex-RNAs in hESC, KMBC, B-cell, PKC, A549, and DKO cells. (B) heatmap of the abundance of the top-three candidate markers in hESC, KMBC, B-cell, PKC, A549, and DKO cells. Each type of cells has three replicates. (C) the scores of the top 9 intracellular smRNAs in H9, NSC, and EC cells. Each type of cell has three replicates. (D) heatmap of the abundance of top-two candidate markers in H9, NSC, and EC cells.

### Validation of stem cell biomarker in diverse datasets and by experiments

For H9 cells, MIR518B and MIR518C were selected as the cell-type markers by our method. For the purpose of validation, datasets in NCBI GEO database for transcriptome studies of miRNAs in hESC cells were collected and the fold change of miR-518b and miR-518c expression levels between hESC and other cell types, including differentiated embryoid bodies derived from H9 cells at 10 days (GSE12229) ^28^, cells at day 4 of H9 differentiation to definitive endoderm (GSE16690) ^29^, isogenic spontaneously differentiating cells from hESC (GSE21722) ^30^, and adult fibroblast cell (HFF-1 cell line) (GSE62501) ^31^. The result is shown in Figure 3(A). Both mir-518b and mir-518c were highly expressed in hESC and have extremely low expression levels in isogenic spontaneously differentiating cells (GSE21722) ^30^ and adult fibroblast cells (GSE62501) ^31^. The fold changes are over 1000 folds. MiR-518b and miR-518c had significantly higher expression levels in H9 than in differentiated cells (GSE12229 ^28^ and GSE16690 ^29^). The fold changes are close to or larger than two. All of these studies agree with the discovery from our system of H9 and adult cells derived from H9 cells – miR-518b and miR-518c are highly expressed in hESC and can work as cell type markers for hESC.

**Figure 3.**
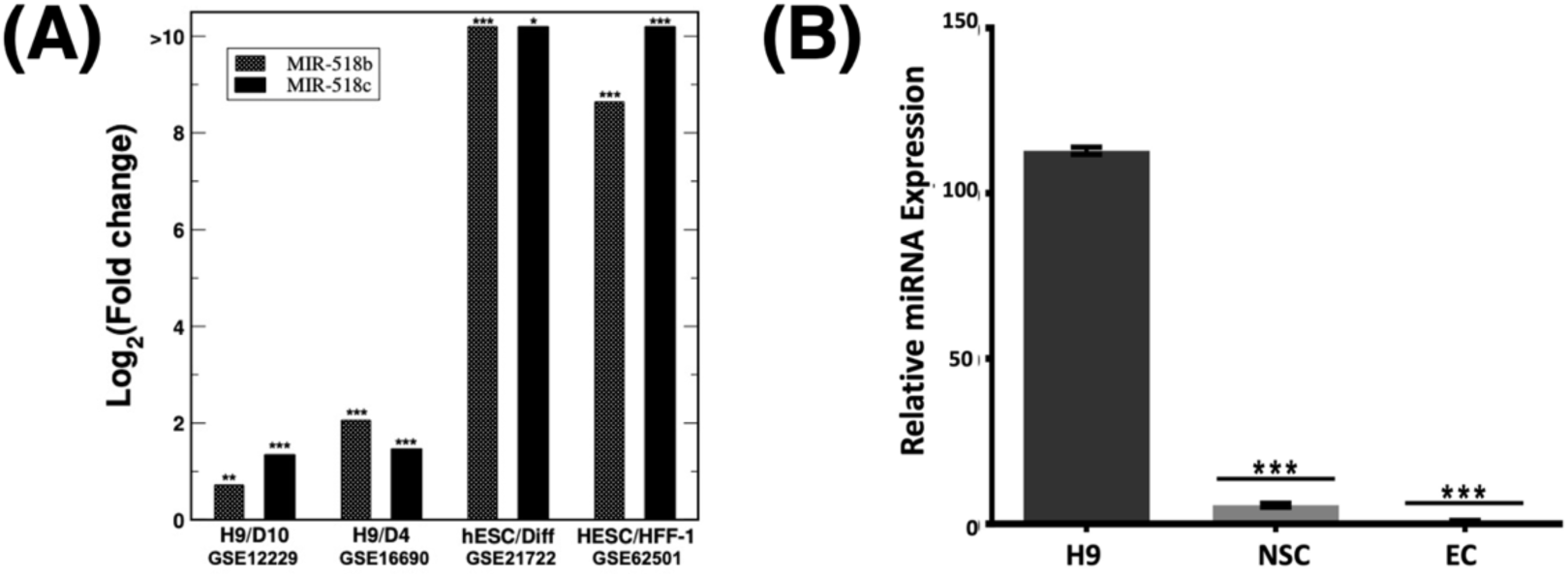
Expression levels of miR-518b and miR-518c in different cells from various datasets and experiments. (A) Fold change of miR-518b and miR-518c expression levels between hESC and other types of cells. * P-value <0.1, ** P-value <0.01, *** P-value <0.0001. (B) qPCR results of miR-518b in H9, NSC, and EC. Data is represented with mean ± standard deviation. *** P-value <0.001

We conduct real-time qPCR to validate if miR-518b is exclusively expressed in H9, but has low expression in NSC and EC. The real-time qPCR product was visualized on Agarose gel. The size of the target was as expected. The DNA was collected and purified for sequencing. The results showed the sequence of the target was the same as expected. The normalized expression levels of miR-518b in H9, NSC, and EC cells are shown in Figure 3(B). In H9, miR-518b was highly expressed, while in NSC and EC, its expression levels are extremely low. Especially, in EC, it even could not detect miR-518b. This result from the small-scale experiment agrees with the discovery of our scoring function on the RNA-seq data.

### Validation of the power of biomarkers for cell-type prediction with a deep-learning approach

To test if discovered markers can be used to identify cell types, we use a simulated data set and neural network classifier to validate discoveries. For the simulation, we use the measured existing RNA-seq datasets as seeds to generate simulated intracellular or extracellular small RNA gene expression profiles based on our RNA-Seq count tables. For the simulated data, 8,991 new RNA-seq data sets were created following the exact distribution of the original data from 9 samples with variation within 0.75 of standard deviation applied, i.e. each has 999 reproduced RNA-seq datasets. A large standard deviation (0.75) was introduced to create very noisy data because the quality of iPSCs may vary depending on the culturing skills ^32^. The total size of the data set was 9000 samples and was separated into two datasets: the training dataset (7,200) and the test dataset (1,800). We applied a neural network classifier and used our discovered maker genes as features to classify the target cell type or non-target cell type.

For the extracellular case, four top ranked biomarker ex-RNAs are features for each cell type, e.g. *MTRNR2L2, TMEM215, MT-TN:Mt_tRNA*, and *SNORD118*, as features to distinguish hESC from the other five cell types. We call this feature set as the marker feature set (MF). For model training, the procedure of three-fold cross-validation was executed 1000 times, and the average value of all areas under the receiver operating characteristic (ROC) curves was used as a criterion to select the optimized parameters. ROC curves, generally displayed as the True Positive Rate (TPR) *versus* the False Positive Rate (FPR), are a popular tool for comparative analysis of computational models and/or diagnostic biomarkers. The area under the ROC curve (ROC-AUC), which varies between 0 and 1, is a measure of discriminate accuracy ^33,34^. A ROC-AUC value close to 1 indicates high TPR and low FPR. After training, ROC-AUC values for optimized models are 0.91, 0.91, 0.90, 0.91, 0.92, and 0.86 for the cell type of hESC, KMBC, B-cell, PKC, A549, and DKO, respectively (Figure 4A). Then, the trained model was applied to the test set. To evaluate the prediction performance, the AUC values for ROC and the area under the precision-recall curve (PR-AUC) values were calculated. The ROC-AUC (PR-AUC) values are 0.97(0.95), 0.94(0.90), 0.96(0.94), 0.99(0.98), 0.98(0.97), and 0.86(0.71), for hESC, KMBC, B-cell, PKC, A549, and DKO, respectively (Figure 4B). This indicates that those selected ex-RNA markers could accurately identify their corresponding cell types.

**Figure 4.**
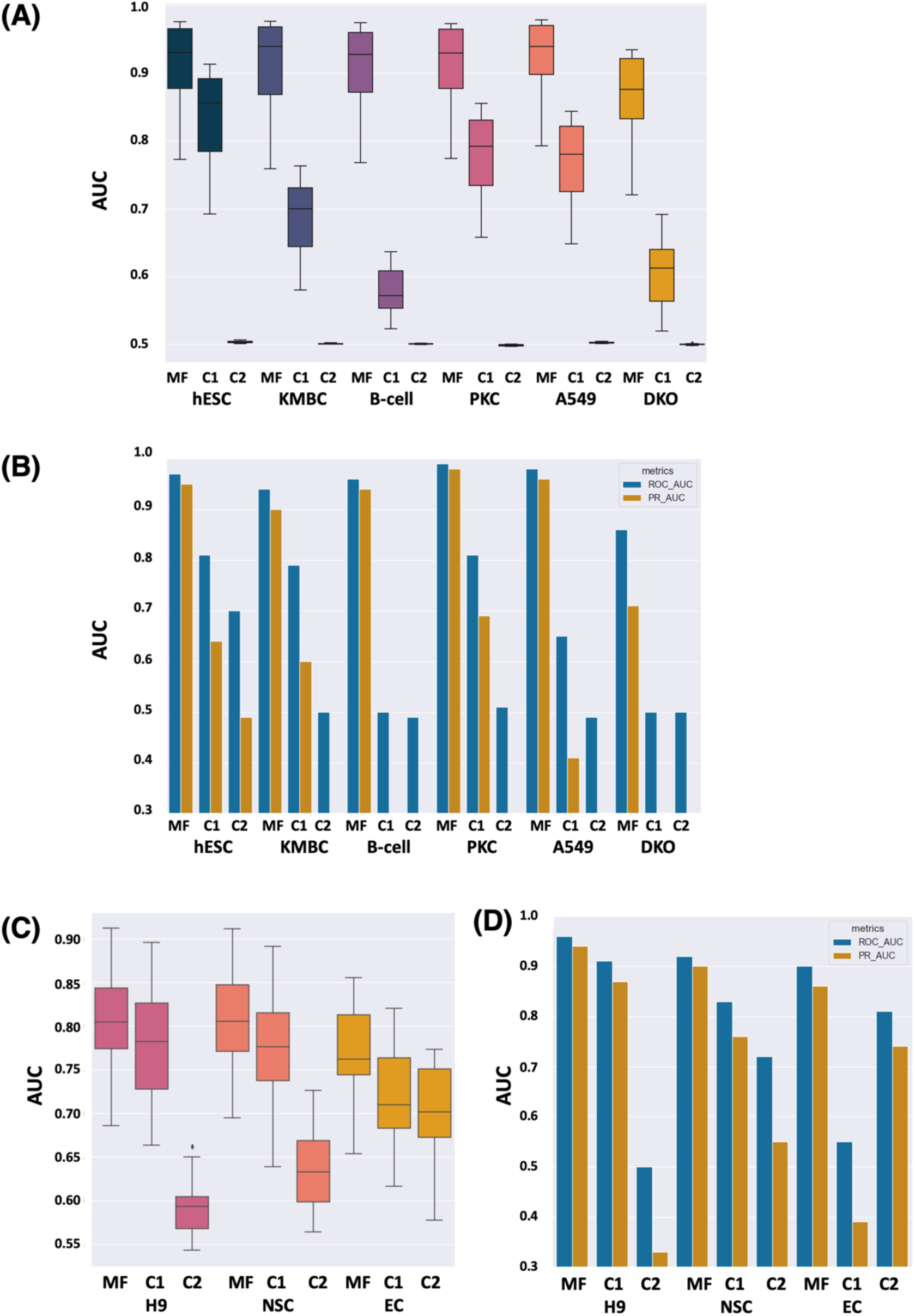
AUC and PR-AUC for training and test datasets. (A) the average values of ROC-AUCs for the training procedures for ex-RNA feature sets for three target cell types. (B) ROC-AUCs and PR-AUCs for the test data for ex-RNA feature sets for three target cell types. (C) the average values of ROC-AUCs for the training procedures for intracellular miRNA feature sets for three target cell types. (D) ROC-AUCs and PR-AUCs for the test data for intracellular miRNA feature sets for three target cell types.

As a control, we picked another two sets of feature ex-RNAs/smRNAs. The first feature set includes ex-RNAs/smRNAs that still have small scores but are higher than the scores of our discovered markers, and the second feature set has the same number of ex-RNAs/smRNAs randomly selected with high scoring function scores. We call these two feature-sets the control set 1 (C1) and control set 2 (C2). A higher score assigned by the scoring function indicates a lower ability to predict cell type for a gene. The same datasets and training/testing procedures were used. After training, average ROC-AUC values for optimized models for C1 feature set are 0.83, 0.79, 0.60, 0.78, 0.77, and 0.60 for the cell type of hESC, KMBC, B-cell, PKC, A549, and DKO, respectively, and C2 feature set are 0.50, 0.50, 0.50, 0.49, 0.50, and 0.59 for the cell type of hESC, KMBC, B-cell, PKC, A549, and DKO, respectively (Figure 4A). With the first control feature set, C1, the final testing ROC-AUC (PR-AUC) values for hESC, KMBC, B-cell, PKC, A549, and DKO are 0.81 (0.64), 0.79 (0.60), 0.5(0.17), 0.81(0.69), 0.65(0.41), and 0.50(0.17), while the second control feature set, C2, lead the ROC-AUC (PR-AUC) values for hESC, KMBC, B-cell, PKC, A549, and DKO to 0.07 (0.49), 0.50 (0.15), 0.49(0.16), 0.51(0.16), 0.49(0.16), and 0.50 (0.17) (Figure 4B). One can see that all AUC values for C1 and C2 features are smaller than those selected ex-RNA markers. The performance of C1 features is better than C2 features; in most cases, C2 features have no ability to predict cell types. After all, ex-RNAs/smRNAs in C1 feature set have relatively small scores and they have a certain level of ability for prediction.

For the intracellular case, we executed the same procedure. During the training procedure, the average ROC-AUC for selected smRNA markers is 0.81, 0.81, and 0.77, for H9, NSC, and EC, respectively, but for C1(C2) feature sets, average ROC-AUCs are 0.78(0.59), 0.78(0.64), and 0.72(0.70), for H9, NSC, and EC, respectively (Figure 3C). ROC-AUCs (PR-AUCs) on the independent test are 0.96 (0.94), 0.92 (0.90), and 0.90 (0.86) for H9, NSC, and EC, respectively (Figure 3D), while ROC-AUCs (PR-AUCs) for other two control feature sets are 0.91 (0.87), 0.83 (0.76), and 0.55 (0.39) for H9, NSC, and EC, for the first feature set, C1, and 0.50 (0.33), 0.72 (0.55), and 0.81 (0.74) for H9, NSC, and EC, for the second random feature set, C2, (Figure 3D). One can get the same conclusion that those selected smRNA markers have better prediction performance than other smRNAs in C1 and C2 feature sets, and smRNAs in the C1 feature set have better performance than randomly selected smRNAs in C2 feature set.

### A deep-learning model to distinguish discovered biomarkers from other non biomarker smRNAs

We developed a machine learning based approach to classify biomarkers with other non-biomarker ex-RNAs/smRNAs. The positive samples in the training set are discovered ex-RNAs/smRNAs identified by the scoring function. We use a multilayer perceptron neural network as a classifier to find if we can distinguish positive biomarkers from other transcripts that cannot be used as biomarkers to decide cell types. Because the number of positives is too small, we simulated more positives using discovered ex-RNAs/smRNAs’ expression levels as seeds. More positives were simulated to make the total number of positives 3,000. The negative set also has 3,000 ex-RNAs/smRNA transcripts that were randomly selected from the original RNA-seq data, excluding our biomarker candidates. The simulated dataset was split into training data set (3,600) and a test dataset (2,400). The model parameters were optimized with training data to prevent information leak. The multilayer perceptron neural network was recursively fitted, and 3-fold cross-validation and grid search were used for the best parameters. With the parameter search, our 2-layer perceptron network had three hidden units on each hidden layer, using ‘idenditi’ activation function, and α=0.3. For ex-RNA biomarkers, the prediction model on the test dataset got ROC-AUCs of 0.98, 0.96, 0.97, 0.92, 0.93 and 0.82 for hESC, KMBC, B-cell, PKC, A549, and DKO cells. The ROC-AUC was 0.93 for H9, 0.76 for NSC, and 0.89 for EC.

## Discussion

For intracellular small RNA-seq data, about 423 miRNAs were detected (>5 reads for one miRNA). Most intracellular miRNAs are constantly expressed in all three types of cells. The most expressed miRNAs in H9 cells are miR-302a,b,c,d, miR-148a, miR-182, miR-183, and miR-93. They are also highly expressed in NSC. There are more than 27,000 transcript accessions (RPM>0.3) for hESC cells reported in the extracellular smRNA dataset obtained from Ref. ^9^. Many tRNAs, snRNAs, and snoRNAs were detected in cell culture media with many reads, such as transcripts from genes *RNU5E-1* and *RNY1*. Out of 27,000 IDs, there are only 63 miRNAs, including miR-302 and miR-3676, and these miRNAs did not have many reads. It has been found that miRNA expression abundance in cells and cell culture media is highly correlated, the extracellular levels of miRNAs are proportional to cell numbers, and the abundance of some miRNAs in cell culture media has a large fluctuation along the differentiation process ^14^. Although there is a correlation between miRNA expression abundance in cells and cell culture media, for example, the miR-302 family, the reads that were detected for miRNAs in the cell culture media are too low to serve as a cell type marker according to the data from Ref. ^9^.

MiRNAs from the same family are often clustered in chromosomal locations^35^. There are two miRNA clusters, miR-302 cluster on chromosome 4 and miR-520 cluster on chromosome 19, that are highly expressed in undifferentiated hESCs ^28^. For intracellular small RNAs, the miR-302 family, miR-367, and miR-373 have been identified as miRNAs specific to embryonic stem cells (ESCs) and iPSCs ^5,36,37^. Intracellular miR-302a and miR-302b were detected in all three types of cells, H9, NSC, and EC, with high expression levels with no significant difference among them. Both miR-367 and miR-373 were highly expressed in H9 and NSC, but had low expression levels in EC. Because there is no significant difference between H9 and NSC, miR-367 and miR-373 could not be used as a biomarker for a system that has both H9 and NSCs. The cluster of miR-520 is composed of 21 miRNAs: miR-518b, miR-518c, miR-519b, miR-519c, miR-519e, miR-520a, miR-520b, miR-520c, miR-520d, miR-520e, miR-520f, miR-520g, etc., and are highly expressed in stem cells in the differentiated bone marrow and embryonic stem cells such as H9 ^38,39^. MiR-518b and miR-518c were selected as the candidate cell-type markers for H9 cells by our method. MiR-518b was found to regulate FOXN1 in the epithelial lineage development in NT2/D1 and H9 hESC^40^. Some fragments of *THBS1* gene’s transcripts were found in ECs and the gene *THBS1* was identified as a candidate marker. The gene *THBS1* has limited expression in the healthy adult, and its gene product is secreted by many cell types in response to injury or specific cytokines ^41^. Usually, THBS1 protein is present transiently in the extracellular matrix but is rapidly internalized for degradation by ECs ^41^.

For extracellular small RNAs, during iPSCs generation and differentiation in mouse, specific ex-RNAs were found in culture media for fibroblasts (miR-214), iPSCs (miR-294, miR-292-3p, miR-323-3p), iPSCs-derived neurons (miR-7-5p) and cardiomyocytes (miR-499a-5p) ^14^. Their abundances had correlations with cell densities. In mESCs, the miR-290-295 cluster accounts for more than 60% of the miRNA population ^42^. In fact, the mouse miR-290-295 cluster is the homolog to human miR-371-373, and the human miR-302 and mouse miR-290-295 clusters share the same seed sequence ^42^. In the dataset for ex-RNAs that we obtained from Ref. ^9^, human miR-302 has the most reads for all miRNAs in hESCs, but compared with other types of small RNA, extracellular miRNAs have very low abundance. Interestingly, mitochondria tRNA fragments, i.e. ex-RNAs from *MT-TN:Mt_tRNA*, have high abundance in the cell culture media of hESC. This type of RNAs, tRNA-derived fragments, are generated by the cleavage of tRNAs. It has been demonstrated that tRNA-derived fragments in extracellular vesicles act as regulatory molecules in various cellular processes and have the potential to be biomarkers in human diseases^43^. SNORD118 was ranked as the fourth top-ranked candidate in all ex-RNAs for hESC cells. Interestingly, SNORD was reported to be discovered in extracellular vesicles released from cells ^44^. It has been discovered that SNORD family was exclusively present in extracellular vesicles from human stem cells ^45^. Some fragments of the transcript of the gene *MTRNR2L2* were detected in the cell medium. It was reported that *MTRNR2L2* and miR-182 are involved in Epigenome regulation for the development of human chronic diseases^46^.

In iPSC-derived regenerative medicine products, it is important to identify the undifferentiated cells. iPSC-biomarkers can increase detection sensitivity. For the development of an automated real-time monitoring system for cell manufacture, a good cell type marker needs to have both good detection sensitivity and specificity. For this goal, we need to find candidate marker genes that have discrete gene expression levels among different cell types, i.e. these marker genes have high expression in one cell type but low/no expression in the other a few cell types. Therefore, some previously discovered candidate marker genes cannot work well. For example, miR-302 family has a high abundance in hESCs, but NSCs also have a high abundance of miR-302. The miRNAs in miR-302 family cannot distinguish hESCs, but NSCs. However, our discovered miR518b and c have a high abundance in H9 cells only. This indicates that using the scoring function that we developed here can rank good cell-type marker candidates on the top of all genes, and their ability to predict cell types is sorted. The prediction performance, i.e. AUC and PR-AUC values for the prediction model used on the independent test dataset, decreased when the abundance of smRNAs that were not ranked on the top by the scoring function was used as markers (Please see the Results Section). We could successfully identify some cell-type markers with our scoring function because, in the system that we are studying, there is a limited number of cell types. In our case, there are only three different cell types. If we can find a good cell-type biomarker, it is also related to the similarity among the different cell types that we are studying. For the extracellular dataset, three types of cells have less similarity than the cell types that we used to get intracellular smNRAs. The discovered biomarker candidates from the extracellular dataset have better performance to predict cell types than the biomarker candidates identified from the intracellular dataset. The number of good biomarker candidates from the extracellular dataset is also larger than that from the intracellular dataset.

## Methods

### Routine cell culture in 2D plates and Differentiation

H9 hESCs (#WA09) were purchased from WiCell Research Institute. Fib-iPSCs were obtained from Human Embryonic Stem Cell Core, Harvard Medical School. Fib-iPSCs were reprogrammed from fibroblasts, by George Q. Daley Lab (Children’s Hospital Boston, MA) and have been well characterized and described in the literature [43]. For 2D cell culture, H9s and iPSCs were cultured in 6-welll plate coated with Matrigel (BD Biosciences, #354277) in Essential 8™ medium (E8, Life Technologies, #A1517001). For differentiation, we followed our published protocols to prepare hPSCs-derived endothelial cells (ECs) and neural stem cells (NSCs) ^25,27,47^.

### RNA sequencing and data preprocessing

Total RNAs for RNA sequencing were extracted from cultured cells with RNeasy mini kit (cat # 74104 QIAGEN) according to the manufacturer’s instruction. Libraries were prepared with TruSeq Stranded mRNA Library Prep Kit and sequenced with Illumina NextSeq 500. 20 million 75 bp paired-end reads were generated for each sample.

Small RNA-seq data analyses were performed as described in Ref. ^48^. After removing the adaptor sequence, the small RNA reads were mapped to the human genome using Bowtie2 program ^49^. Normalization was performed using the total numbers of readings.

### RNA extraction and Real-time qRT-PCR

Total RNA was extracted from cultured cells using Trizol reagent (Invitrogen, USA). First-strand cDNA was synthesized from total RNA using M-MLV reverse transcriptase kit with the stem-loop RT primer of miRNAs and using Multiscribe Reverse Transcriptase kit (Applied Biosystems) with random primers according to the manufacturer’s protocol. Real-time qRT-PCR was performed for miRNAs and U6 using the SYBR Green master mix (Bimake). The relative expression levels of miRNAs were calculated using the 2^−ΔΔCt^ analysis method by normalization to the expression level of U6. The primers used in these experiments are described in Supplementary Table S1.

### Ex-RNA data

Transcriptomic data of ex-RNAs in the medium for the cell culture system were obtained from Ref. ^9^. Two types of human cells, hESC and KMBC, were reported. We selected the data generated from the same lab (Lab5) with the same kit (ExoRNeasy). All transcriptomic data for these two cell types were processed and normalized by Ref. ^9^. Transcriptomic data of ex-RNAs in peripheral blood B lymphocytes were obtained from GSE74759 in GEO database (three samples: GSM1931811, GSM1931815, and GSM1931816) ^20^. Transcriptomic data of ex-RNAs of Epidermal primary keratinocytes (PKCs) came from GSE106453 in GEO database (three replicates: GSM2837765, GSM2837769, and GSM2837773) ^21^. Data for lung cancer cell lines (A549) were obtained from GSE106277 (three replicates: GSM2834518, GSM2834519, and GSM2834520) ^22^. Transcriptomic data of ex-RNAs of DKO cell line of colorectal cancer (DKO) were collected from In GSE125905 (three samples: GSM3584527, GSM358452, and GSM3584530) ^23^. All RNA-seq data were trimmed with Cutadapt (v4.1), and reads were mapped human reference genome, GRCh38, with HISAT2^50^. Reads in each gene or transcript were counted by HTSeq^51^. RPM of each gene’s read count was used to make the gene expression levels in different cell types comparable.

### Intracellular smRNA transcriptomic datasets

Dataset 1: H9 and differentiated embryoid bodies derived from H9 cells at 10 days (GSE12229) ^28^, Dataset 2: H9 and cells at day 4 of H9 differentiation to definitive endoderm (GSE16690) ^29^, Dataset 3: hESC and isogenic spontaneously differentiating cells from hESC (GSE21722) ^30^, and Dataset 4: hESC and adult fibroblast cell (HFF-1 cell line) (GSE62501) ^31^ were collected from NCBI GEO database. Microarray data were preprocessed, including background correction and normalization, with *rma* function from R *affy* package ^52^. By averaging the expression of probes corresponding to the same gene symbol.

### Deep learning-based prediction

The multi-layer perceptron network was built with scikit-learn (v.1.0.2), one hidden layer or two hidden layers with three hidden units; the activation function, optimal training steps, and iterations were found by grid search with three-fold cross-validation. Data were split into test data and training data. During the training procedure, three-fold cross-validation was used to find the best activation function, learning step, and optimum hidden layer structures.

### Synthetic data by simulation

Limited by the amount of data available, we use synthetic data to test our model. Synthetic data was created with ‘NumPy’ (v 1.22.0) by adding random noise on the count matrix by following a normal distribution with a standard deviation of 0.75. Normalization by count per million was done by ‘bioinfokit’. To test if discovered biomarkers can be used as features to distinguish the target cell type from other cell types, more sets of cell transcriptomes were synthesized (8,991 new data sets). To test if discovered biomarkers can be distinguished from other transcriptions, new biomarker-like transcripts were synthesized.

## Supporting information

Primer sequences

## Author contributions

Experimental design and model construction, YS, YL, and CZ; Cell culture and RNA extraction, HL, WY, and LH; data analysis and interpretation, AC, RS, and CZ; supervision and editing the manuscript, CZ and YL. All authors have read and agreed to the published version of the manuscript.

## Data availability

All the data that support the findings of this study are available within the paper and its supplementary information files. The small-RNA-seq data are available at NCBI GEO (GSE206925).

